# Structural distortions induced by Kinase Inhibiting RNase Attenuator (KIRA) compounds prevent the formation of face-to-face dimers of Inositol Requiring Enzyme 1α

**DOI:** 10.1101/744904

**Authors:** Antonio Carlesso, Chetan Chintha, Adrienne M. Gorman, Afshin Samali, Leif A. Eriksson

## Abstract

Inositol-Requiring Enzyme 1α (IRE1α) is a transmembrane dual kinase/ribonuclease protein involved in propagation of the unfolded protein response (UPR). IRE1α is currently explored as a potential drug target due to growing evidence of its role in variety of disease conditions. Upon activation, IRE1 cleaves X-box Binding Protein 1 (XBP1) mRNA through its RNase domain. Small molecules targeting the kinase site are known to either increase or decrease RNase activity, but the allosteric relationship between the kinase and RNase domains of IRE1α is poorly understood. Subsets of IRE1 kinase inhibitors (known as “KIRA” compounds) bind to the ATP-binding site and allosterically impede the RNase activity. KIRA compounds are able to regulate the RNase activity by stabilizing monomeric form of IRE1α.

In the present work, computational analysis, protein-protein and protein-ligand docking studies, and molecular dynamics simulations were applied to different IRE1 dimer systems to provide structural insights into the perturbation of IRE1 dimers by small molecules kinase inhibitors that regulate the RNase activity. By analyzing structural deviations, energetic components and number of hydrogen bonds in the interface region, we propose that the KIRA inhibitors act at an early stage of IRE1 activation by interfering with IRE1 face-to-face dimer formation, thus disabling the activation of the RNase domain. The work sheds light on the mechanism of action of KIRA compounds and may assist in development of further compounds in e.g. cancer therapeutics. The work also provides information on the sequence of events and protein-protein interactions initiating the unfolded protein response.

**Non-technical Summary:** The unfolded protein response is a protective feedback mechanism whereby cells regulate high levels of misfolded proteins in the endoplasmic reticulum. Due to its significance in cell survival, the UPR has become an interesting target in cancer therapy. A key pathway of the UPR is initiated by the activation of inositol requiring enzyme 1α (IRE1α), which must first dimerise in order to mediate the stress signal. Different inhibitors have been proposed in order to block the UPR at the level of IRE1α. We here unveil, through detailed computational studies, the mode of action of a set of IRE1α inhibitors targeting the kinase domain, which in turn helps us to further understand the mechanism of activation and progression of the UPR.

## 1. Introduction

The accumulation of misfolded proteins in the endoplasmic reticulum (ER) triggers an evolutionarily conserved intracellular signaling pathway called the unfolded protein response (UPR)(1). The UPR is mainly an adaptive response to re-establish ER proteostasis through the signaling of three transmembrane proteins, inositol-requiring protein 1α (IRE1α, hereafter IRE1), protein kinase R (PKR)-like ER kinase (PERK), and activating transcription factor 6 (ATF6)(2). IRE1 represents the most evolutionarily conserved branch of the UPR: it is a multidomain protein with an N-terminal luminal domain and cytosolic kinase and RNase domains connected by a transmembrane linker(2). Upon activation, IRE1 dimerizes, trans-autophosphorylates and oligomerizes, thereby activating the IRE1 RNase domain which is able to remove a 26 nucleotide intron from X-Box Binding Protein 1 (XBP1) mRNA. Upon re-ligation by RNA-splicing ligase RtcB, spliced XBP1 (XBP1s) is formed(2). XBP1s protein is a potent transcription factor and its target genes in turn permit the ER to adapt to stress conditions(2). There are several models that explain how ER stress is sensed by IRE1(3). One of the established models links its trans-autophosphorylation with face-to-face dimer orientation as the first step of IRE1 activation (Figure 1A), and subsequent transition to back-to-back dimer (Figure 1B) and larger oligomeric structure formation to stimulate the RNase domain activity(3).

**Figure 1.**
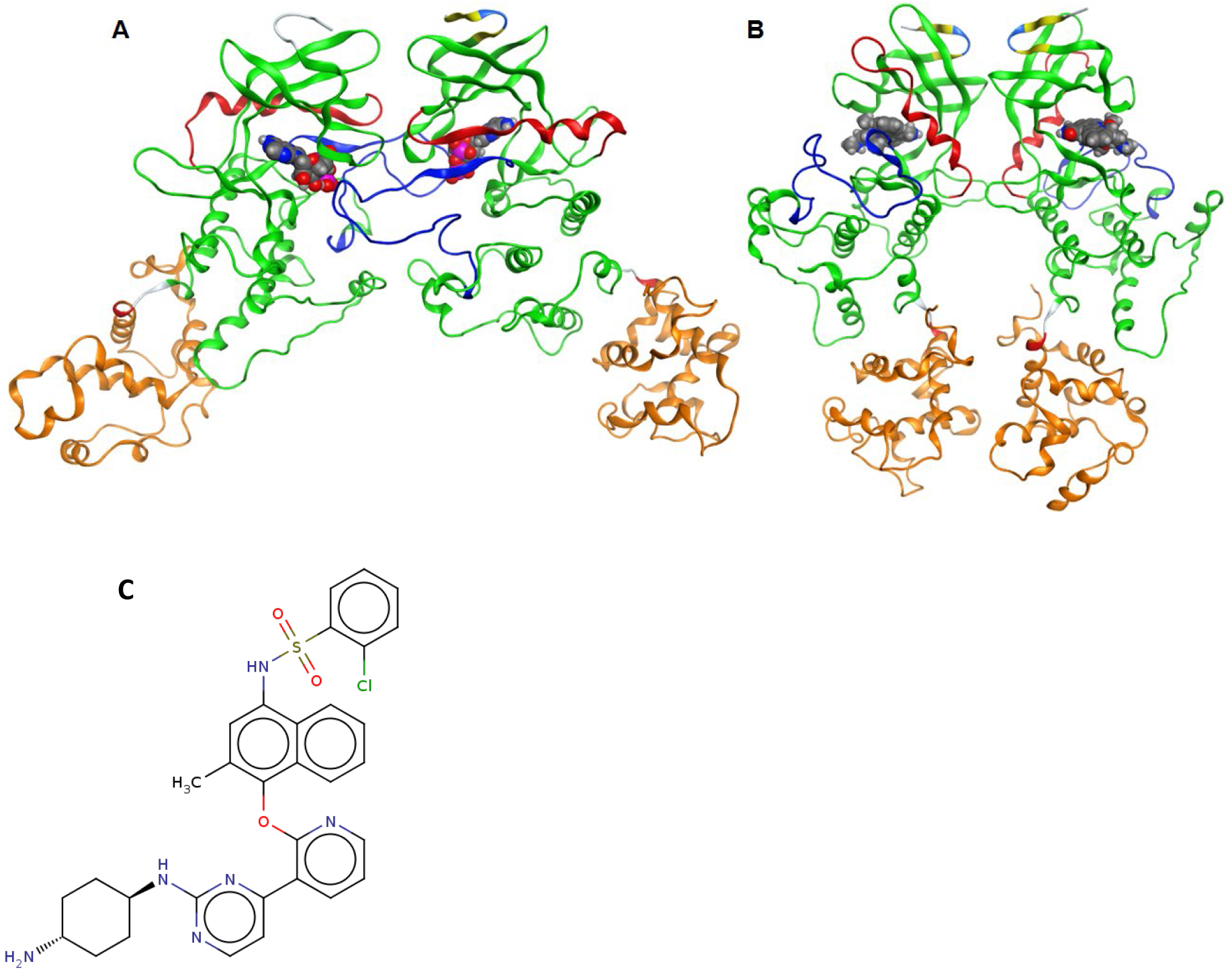
Ribbon diagram representing the structure of the IRE1 kinase and RNase of (A) the face-to-face dimer (PDB 3P23) and (B) back-to-back arrangement (PDB 4YZC). The kinase domain is shown in green (residues 571-832), the helix-αC in red (residues 603-623), the activation segment in blue (residues: 711-741) and the RNase domain in orange (residues 837-963). ADP (A) and staurosporine (B) are highlighted in space-filling models to indicate the kinase binding sites. (C) 2D molecular representation of KIRA.

IRE1 signaling is implicated in the etiology of various diseases(2, 4), including cancer where tumor cells activate the UPR to avoid apoptosis and survive(5). Thus, IRE1 has been the recent focus of several drug discovery projects in cancer research(6). Current IRE1 modulators can be categorized as (i) direct IRE1 RNase inhibitors(6, 7), (ii) ATP-competitive kinase inhibitors that activate RNase(8) and (iii) ATP-competitive kinase inhibitors that decrease RNase activity(8, 9), the latter also known as ‘kinase inhibiting RNase attenuators’ (KIRAs). Optimized KIRA compounds include KIRA6(10), KIRA7(4) and KIRA8(11). KIRA6 and KIRA7 possess an imidazopyrazine scaffold(10), whereas KIRA8 is a sulfonamide compound with high selectivity(11).

The observation that KIRAs allosterically inhibit IRE1 RNase domain was confirmed by a competitive *in-vitro* assay where ATP-competitive RNase activators were shown to completely restore the RNase activity in presence of KIRA(8). The model proposed by Feldman *et al*. speculates that KIRAs stabilize the DFG-out kinase domain conformation and helix-αC displacement that makes it incompatible with back-to-back dimer formation, thereby leading to kinase and RNase inhibition(8).

Herein, we propose an alternative, and we think more likely, model to rationalize the link between kinase binding and RNase domain inhibition by the KIRA compounds based on molecular level analyses of structures and dynamics of IRE1 in complex with KIRA compound **16** (Figure 1C); numbering from the original paper(9) (hereafter referred to as KIRA).

Using protein-protein docking(12), protein-ligand docking(13) and molecular dynamics (MD) simulations(13) we investigate structure and dynamics of the two relevant IRE1 dimer structures (Figure 1), face-to-face and back-to-back, critical for trans-autophosphorylation and RNase activity,(3) respectively. Using models in the presence and absence of KIRA bound to the kinase pocket, we propose a mechanistically-based model of how KIRAs inhibit the RNase activity of IRE1 by allosteric interaction with the kinase domain.

## 2. Methods

### 2.1. Selection and preparation of IRE1 crystal structure

IRE1 crystal structures with PDB codes 4U6R (KIRA-bound monomer), 3P23 (ADP bound face-to-face dimer) and 4YZC (staurosporine bound back-to-back dimer) were prepared using Schrödinger protein preparation wizard(14). Using the 3P23 PDB structure, the face-to-face dimer was obtained by deleting Chain C and D and generating missing loops using Prime(15). Hydrogen atoms were added and the protonation and tautomeric states of Asp, Glu, Arg, Lys and His were adjusted to match a pH of 7.4. Possible orientations of Asn and Gln residues were generated. Finally, the IRE1 dimer and monomer structures were subjected to geometry refinements using the OPLS3 force field(16) in restrained minimizations.

### 2.2. Protein-Protein Docking

A protein-protein docking analysis was performed to understand the structural basis of IRE1 recognition and regulation mediated by KIRA. Initially, five different protein-protein docking programs were chosen: SwarmDock(17), ZDOCK(18), HsymDock(19), PatchDock(20) and ClusPro(21). The ability of the programs to reproduce the native IRE1 dimers (PDB code 4YZC for back-to-back and 3P23 for face-to-face) was checked. The best performing program, SwarmDock(17), was then used for predicting the structures of KIRA-bound dimers using the 4U6R PDB monomer structure. A schematic representation of the method used is shown in Figure 2.

**Figure 2.**
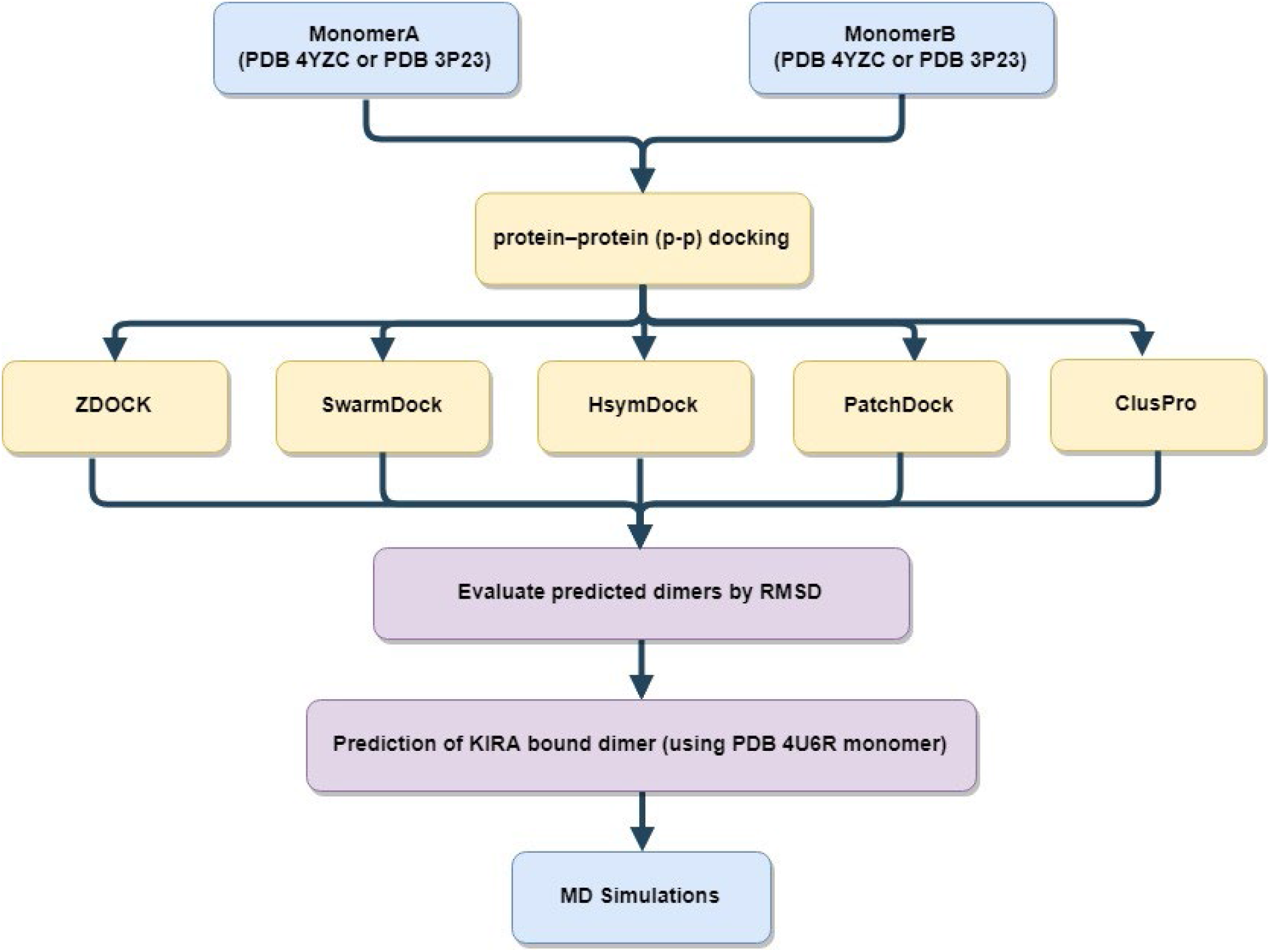
Schematic representation of protein-protein docking scheme used for predicting KIRA-bound dimer structures.

### 2.3. KIRA preparation for docking studies

The co-crystallized ligand from PDB 4U6R as shown in Figure 1C was extracted from the PDB structure and used for docking studies. KIRA is displayed in Figure 1C. KIRA was prepared using LigPrep(22) in the Schrödinger suite(23). The OPLS3 force field(16) was used for KIRA preparation steps and possible protonation and ionization states were assigned using Ionizer at pH 7.4.

### 2.4. Molecular docking of KIRA

KIRA was docked in the kinase pocket of the IRE1 dimer back-to-back (PDB code: 4YZC) and face-to-face (PDB code: 3P23) structures, using the Glide program(24) in Schrödinger(23) with the receptor grids prepared using the OPLS3 force field(16). The molecule was docked in the kinase domain of both monomers of each dimer. IRE1 dimers in apo form were obtained by deleting the small organic molecules and ions present in the crystal structures, from the kinase active site (*i.e*., ADP and Mg^2+^ in the 3P23 PDB structure and staurosporine in the 4YZC PDB structure, respectively). Two grid centers per dimer (*i.e*., one per each kinase active site) were prepared, each cubic grid with a side length of 20 Å. The grid center was set at the centroid of Lys599, a residue crucial for the kinase activity. XP (Extra Precision) docking mode and flexible ligand sampling were employed in the docking procedure. All other parameters were set to default values.

### 2.5. Molecular Dynamics Simulations

The stability of the native crystallographic dimer structures of the IRE1 in back-to-back (PDB 4YZC) and face-to-face (PDB 3P23) conformer were compared with the predicted KIRA-bound dimer forms, based on 300 ns MD simulations. For KIRA bound dimers we used both complexes with KIRA docked in the kinase active site of the dimer structures 4YZC and 3P23, and dimer structures generated from KIRA-co-crystallized IRE1 monomers.

For the MD simulations, the following steps were followed:

a. Systems preparation: Systems include the experimental IRE1 dimer structures (PDB 4YZC, 3P23), predicted dimers (from KIRA bound monomer) (section 2.2), and KIRA-docked dimer forms (section 2.4). The systems were prepared separately as discussed in section 2.1.
b. Ligand parameterization: ligands (ADP, staurosporine and KIRA) were parametrized with GAFF as implemented in Ambertools2018 by using the Antechamber interface tool(25). AM1-BCC atomic point charges(26) were calculated using Antechamber(27).
c. Molecular dynamics simulation protocol: MD simulations were performed using the GROMACS 5.1 package(28), with the AMBER14SB force field for the protein(29). The systems were explicitly solvated using cubic water boxes with cell borders placed at least 10 Å away from the protein or ligand atoms using TIP3P water(30) under periodic boundary conditions. The systems were first neutralized and Na^+^/Cl^−^ counter ions were added to give a physiological salt concentration of 0.154 M. All simulation runs consisted of energy minimization until the force was less than 1000 kJ mol^−1^ nm^−1^, 200 ps under NVT conditions subjected to position restrained equilibration on the heavy atoms of IRE1, 200 ps equilibration and 300 ns of classical molecular dynamics simulation under NPT conditions. The simulations were run in triplicate (referred to as Replica 1, 2 and 3). In all simulations, the temperature was kept at 300 K by the velocity rescaling thermostat(31) with a coupling constant of 0.1 ps, and pressure at 1.01325 bar using the Parrinello-Rahman barostat(32) with a coupling time of 5.0 ps, excluding NVT pre-simulation steps. Constraints were applied on all bonds using the LINCS algorithm(33). The leap-frog algorithm(34) was employed in the simulations, with integration timesteps of 2 fs.

The structural deviations during the MD simulation were analyzed using RMSD, number of distinct hydrogen bonds and energy terms such as electrostatic (Ele) and van der Waals (vdW) interactions using built-in tools in the GROMACS 5.1 package(28). For analyzing the dimer interface RMSD, an index file was created with specific residues. The dimer interface was defined as any pair of Cα atoms from one monomer within 10 Å of the other in the face-to-face and back-to-back dimers(35, 36).

### 2.6. Data availability

All simulation protocols are provided as tarballs (.tar.gz) freely accessible at zenodo.org as DOI: 10.5281/zenodo.3368654.

There are eighteen tarball (**.tar.gz**) files, three replicas for each of the sytems investigated. The contents of each tarball is as follows:

1. a source PDB (**.pdb**) file
2. leap.log - commands used to create the .prmtop and .inpcrd files
3. Two AMBER parameter/topology (**.prmtop**) and an AMBER coordinate (**.inpcrd**) file
4. **.mdp file** used for performing all the minimisation, relaxation, equilibration and production run steps
5. Executable **script (*i.e*. job009)** that was used to perform the production run
6. trajectory (**.xtc**) files for each independent MD simulations

## 3. Results and discussion

### 3.1. Protein-Ligand Docking analysis

KIRA bound IRE1 dimer crystal structures are not available. To predict the KIRA binding in dimers, the compound was docked in the IRE1 back-to-back (PDB code: 4YZC) and face-to-face (3P23) structures. Visual inspection of the docked poses revealed a different binding mode of KIRA compared to the 4U6R PDB structure (Figure S1). In particular, in the face-to-face dimer KIRA binding is mainly stabilized by electrostatic interaction with Asp711, Asp688 and Lys690 (Fig S2A and B) and in the back-to-back dimer with Glu651 and Cys645 (Fig S2C and D) while the co-crystallized KIRA interacts mainly with Lys599, Glu651, Cys645, Phe712 and Ile642 as reported in previous studies(9, 37).

### 3.2. Protein-Protein Docking analysis

To address the questions regarding the effect of KIRA binding on the IRE1 dimer formation, and if KIRA is able to structurally interfere with either the face-to-face or back-to-back dimer form, or both, we assessed if the KIRA-bound monomer structure (PDB code: 4U6R) is capable of forming dimer structures.

In order to identify an appropriate protein-protein docking program, a series of docking experiments were performed using five freely available programs. Starting from the known crystallographic dimer structures of IRE1 in back-to-back (PDB code: 4YZC) and face-to-face (3P23) conformers, we split the crystal structure into monomers and tried to reproduce the dimer complexes with the programs. The docking results are shown in Table 1 and S1.

**Table 1.**
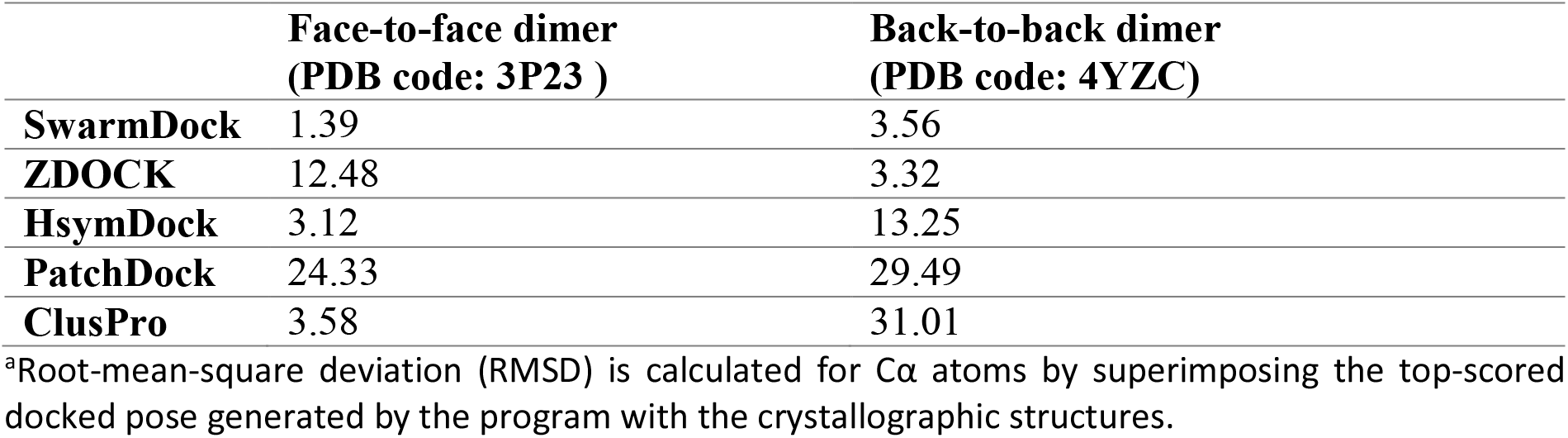
RMSD^a^ results of five different protein–protein docking approaches to reproduce the known IRE1 dimer complexes.

|  | Face-to-face dimer<br>(PDB code: 3P23 ) | Back-to-back dimer<br>(PDB code: 4YZC) |
| --- | --- | --- |
| **SwarmDock** | 1.39 | 3.56 |
| **ZDOCK** | 12.48 | 3.32 |
| **HsymDock** | 3.12 | 13.25 |
| **PatchDock** | 24.33 | 29.49 |
| **ClusPro** | 3.58 | 31.01 |

<sup>a</sup>Root-mean-square deviation (RMSD) is calculated for Cα atoms by superimposing the top-scored docked pose generated by the program with the crystallographic structures.

Evaluating the RMSD over the five top-scored docked poses for all programs studied here (Table S1), we note that the top-scored docking pose in almost all cases is also the one with lowest RMSD. Of the five docking programs tested in the current study, SwarmDock was able to reproduce the native crystallographic back-to-back (PDB code: 4YZC) and face-to-face (PDB code: 3P23) dimers structures of the IRE1 to an RMSD of 3.56 and 1.39 Å, respectively, as shown in Figure 3. SwarmDock was thus also used for predicting dimers of the KIRA-bound monomer structures (PDB 4U6R) in the back-to-back and face-to-face orientations.

**Figure 3.**
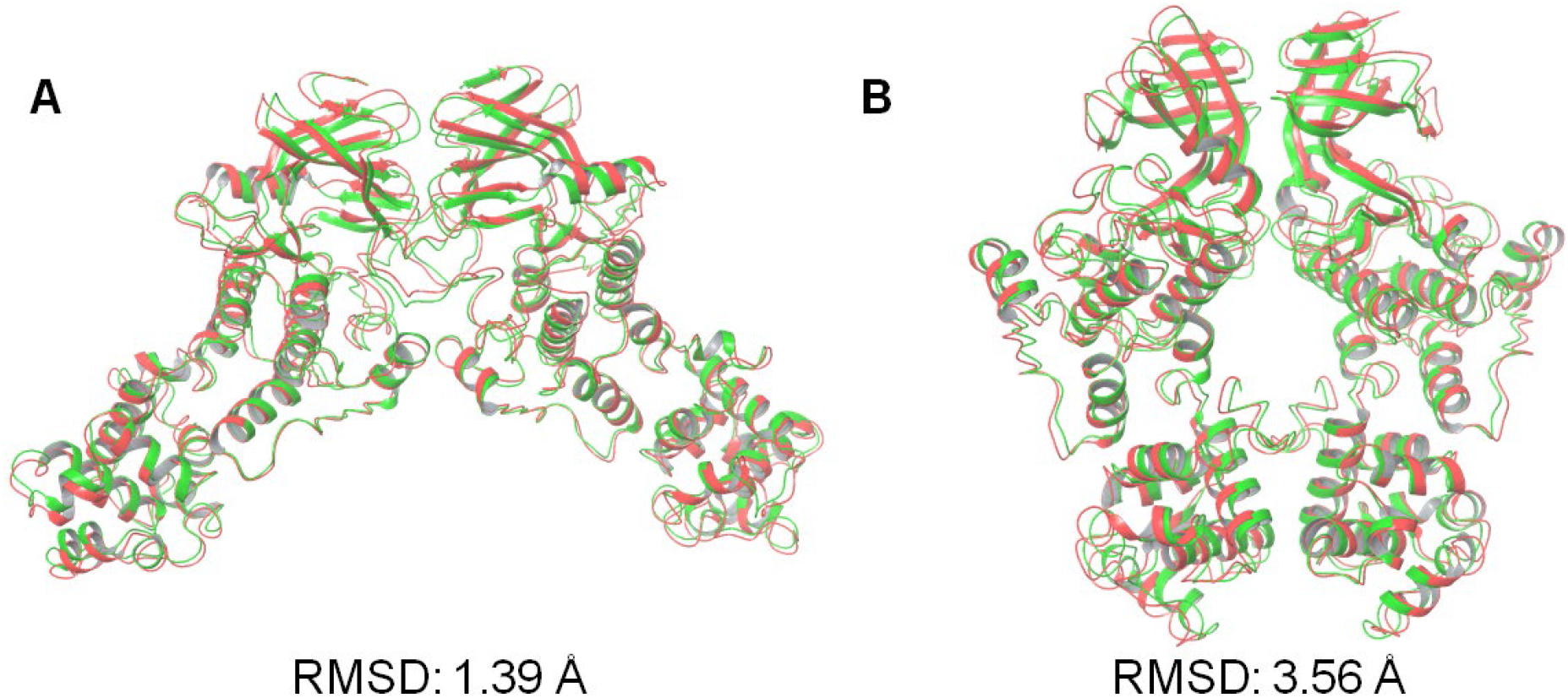
Superposition of the predicted best-scoring poses predicted by SwarmDock (green), onto the crystallographic dimer structures (red) of the IRE1 in (A) face-to-face (PDB code: 3P23) and (B) back-to-back (PDB code: 4YZC) conformations. RMSD values based on the positions of the Cα atoms relative to the crystallographic structures.

We first evaluated steric clashes at the interchain region by superposing the monomer of the 4U6R PDB structure on each monomer of the native crystallographic structures of the IRE1 back-to-back and face-to-face dimers (Figure S3). The KIRA-bound dimer forms produce several steric clashes at interchain level, especially at the helix-αC and the activation segment in KIRA-bound face-to-face dimer (Figure S3). We note that the backbone of Glu604 and the carboxylate group of Glu735 create steric clashes with the guanidino group of Arg600 and the carboxylate of Glu735, located in the helix-αC and the activation segment, respectively (Figure S3). In the back-to-back dimer the guanidine groups of Arg627 and Arg905 create potential steric clashes with the guanidine groups of Arg627 and Arg905 located in the other monomer of the symmetric complex (Figure S3).

Results from the protein-protein docking calculations of the KIRA-bound dimer forms performed using SwarmDock are shown in Figure 4. The best docked results were analyzed further through MD simulations.

**Figure 4.**
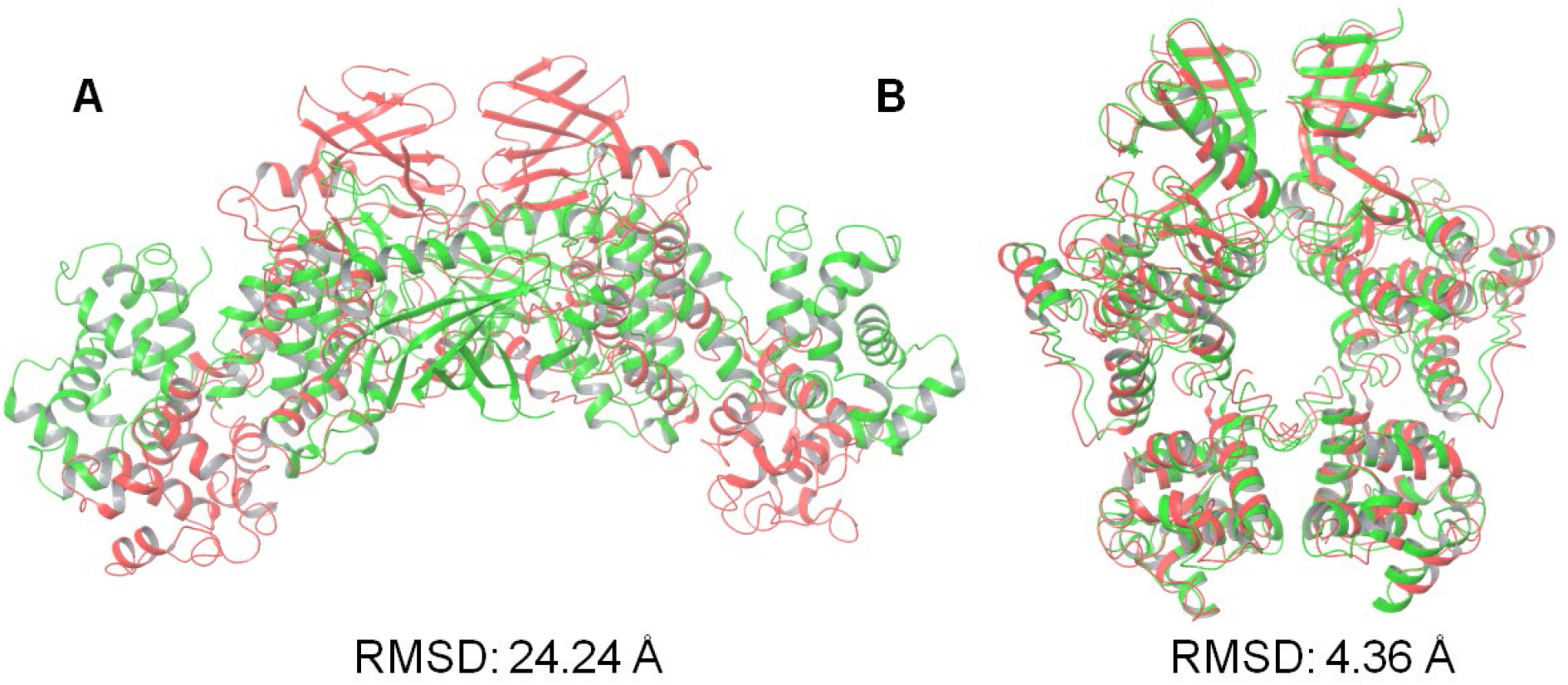
Illustration of the protein-protein docking results of KIRA containing IRE1 monomers (PDB code: 4U6R). Values shown are the RMSD in angstrom in Å of the positions of the Cα atoms of the best-scoring docked pose (green) against the native IRE1 dimer structure in (A) face-to-face (PDB code: 3P23) and (B) back-to-back (PDB code: 4YZC) conformation (red).

The best docked pose (in terms of Cα atom RMSD) generated for the face-to-face dimer shown in Fig. 4A is very far from the experimental one, with an RMSD of 24.24 Å. No structures with better RMSD were found within the 5 top scoring docking poses (RMSD 24.24, 25.29, 34.11, 27.49 and 25.40 Å, respectively). Since the program is shown to successfully predict the crystallographic dimer forms, the high RMSD values are not likely a result of bad sampling. Rather, the KIRA induced conformational changes have rendered the system incapable of appropriate dimer formation.

Comparison of the best back-to-back docking pose obtained using the 4U6R PDB structure and the crystal structures of the back-to-back dimer was also performed (Figure 4B). The RMSD analysis for the best docking pose gives a value near 4 Å, in line with the RMSD value identified when using the native monomer structure (Table 1).

### 3.3. MD simulations analysis: influence of KIRA on the face-to-face dimer

Three different IRE1 face-to-face dimers were explored further, namely the native crystal dimer structure (PDB code: 3P23), the native structure (PDB 3P23) with KIRA docked in the active sites, and the protein-protein docked pose of PDB 4U6R in the face-to-face dimer. The stability of each of the three systems was analyzed during three MD replicas, each replica being 300 ns duration.

To assess the structural stabilities, RMSD values of each dimer and of the dimer interfaces were calculated (Figure 5). For each replica, the calculations were done by considering the structures present in the minimized, equilibrated systems as the reference points. RMSD values were analyzed as the functions of simulation time. The three replicas for the native face-to-face crystal dimer (PDB code: 3P23) reveal that the IRE1 dimer is stable as evidenced by low and relatively constant RMSD values of the three independent trajectories (Figures 5A and D). In contrast, the KIRA-docked face-to-face dimer and the predicted PDB 4U6R dimer reach higher values of RMSD (Figures 5B, C, D and E) indicating that these dimers explore more distorted complexes compared to the native IRE1 face-to-face dimer. Conformations with large interface RMSD were visually examined and by comparing the distance of the center of mass (COM) of each RNase domain in each dimer with the native face-to-face dimer we analyze the impact of KIRA on the stability of the system (Figure S4). The COM distance of the RNase domain in the KIRA docked in PDB 3P23 dimer and in the protein-protein docked pose of PDB 4U6R in face-to-face dimer form are significantly higher compared to native IRE1 dimer demonstrating the impact of KIRA on the stability of the system (Figure S4).

**Figure 5.**
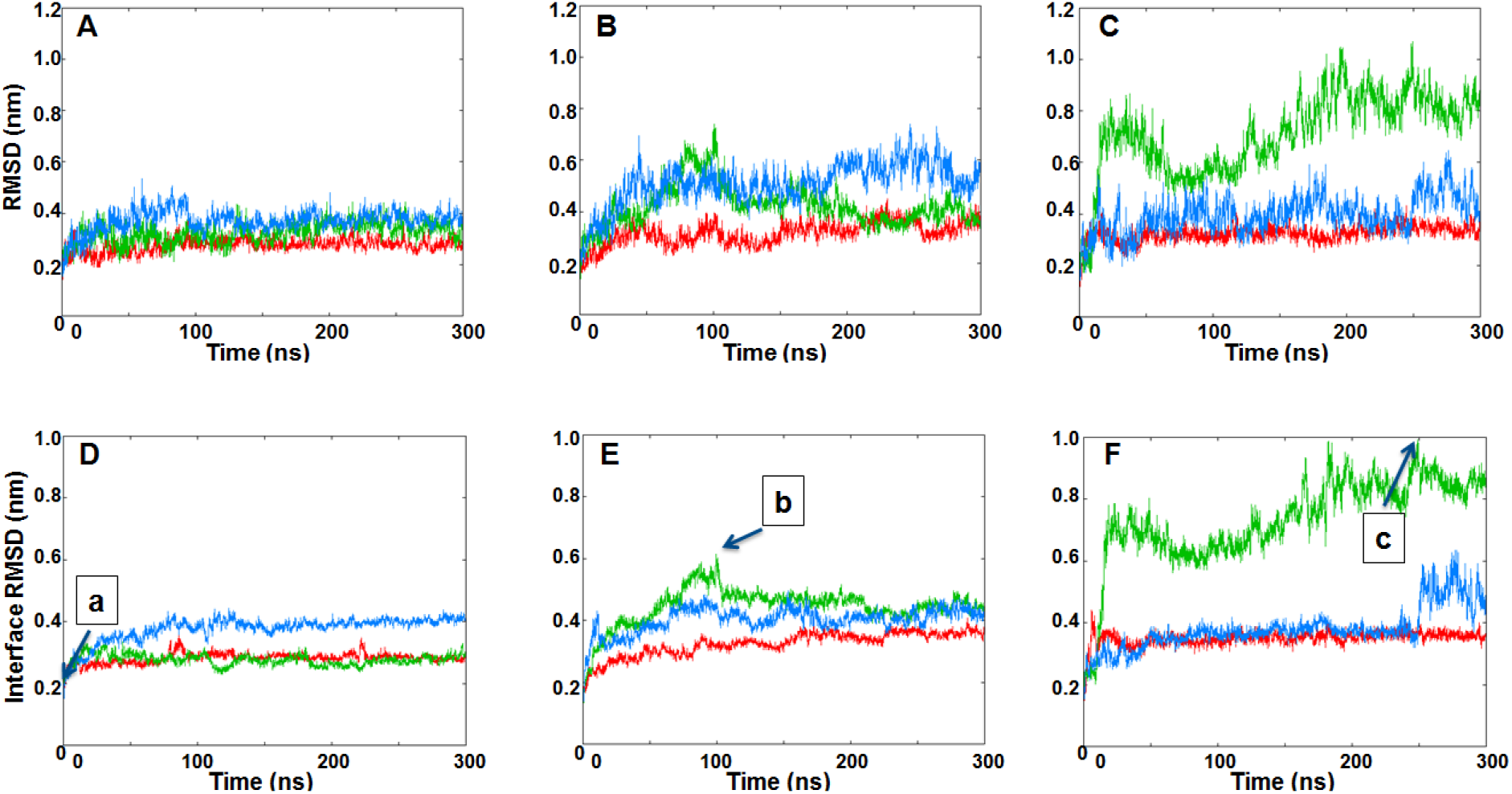
RMSDs of IRE1 face-to-face dimer Cα atoms during the three MD simulation replicas of (A) Native face-to-face crystal dimer structure (PDB code: 3P23), (B) KIRA docked in PDB 3P23 dimer, (C) protein-protein docked pose of PDB 4U6R in face-to-face dimer form. Interface RMSDs of IRE1 face-to-face dimer Cα atoms during the three MD simulation replicas of (D) Native face-to-face crystal dimer structure (PDB code: 3P23), (E) KIRA docked in PDB 3P23 dimer, (F) protein-protein docked pose of PDB 4U6R in face-to-face dimer form. Replicas 1, 2, and 3 are represented in red, green and blue, respectively. Individual frames of the MD simulations labeled a, b and c are shown in Figure S4.

To further analyze the system deviation, the interaction energy and the number of H-bonds between the monomers were computed (Figure 6). Interaction energy analysis between the monomers reveals a smaller energetic stabilization of the KIRA-docked face-to-face dimer and the protein-protein docked pose of PDB 4U6R in face-to-face dimer form, compared to the native face-to-face crystal dimer (PDB code: 3P23) (Figure 6). The same trend was confirmed by the H-bonds analysis, with a higher number of H-bonds occurring between the native IRE1 face-to-face dimer compared to protein-protein docked pose and KIRA-docked face-to-face dimer (Figure 6). The overall analysis confirms our initial interpretation of the different stabilities of the three IRE1 face-to-face dimers investigated, with the native one being the more stable, indirectly reflecting the impact of KIRA on the stabilization of the IRE1 face-to-face dimer.

**Figure 6.**
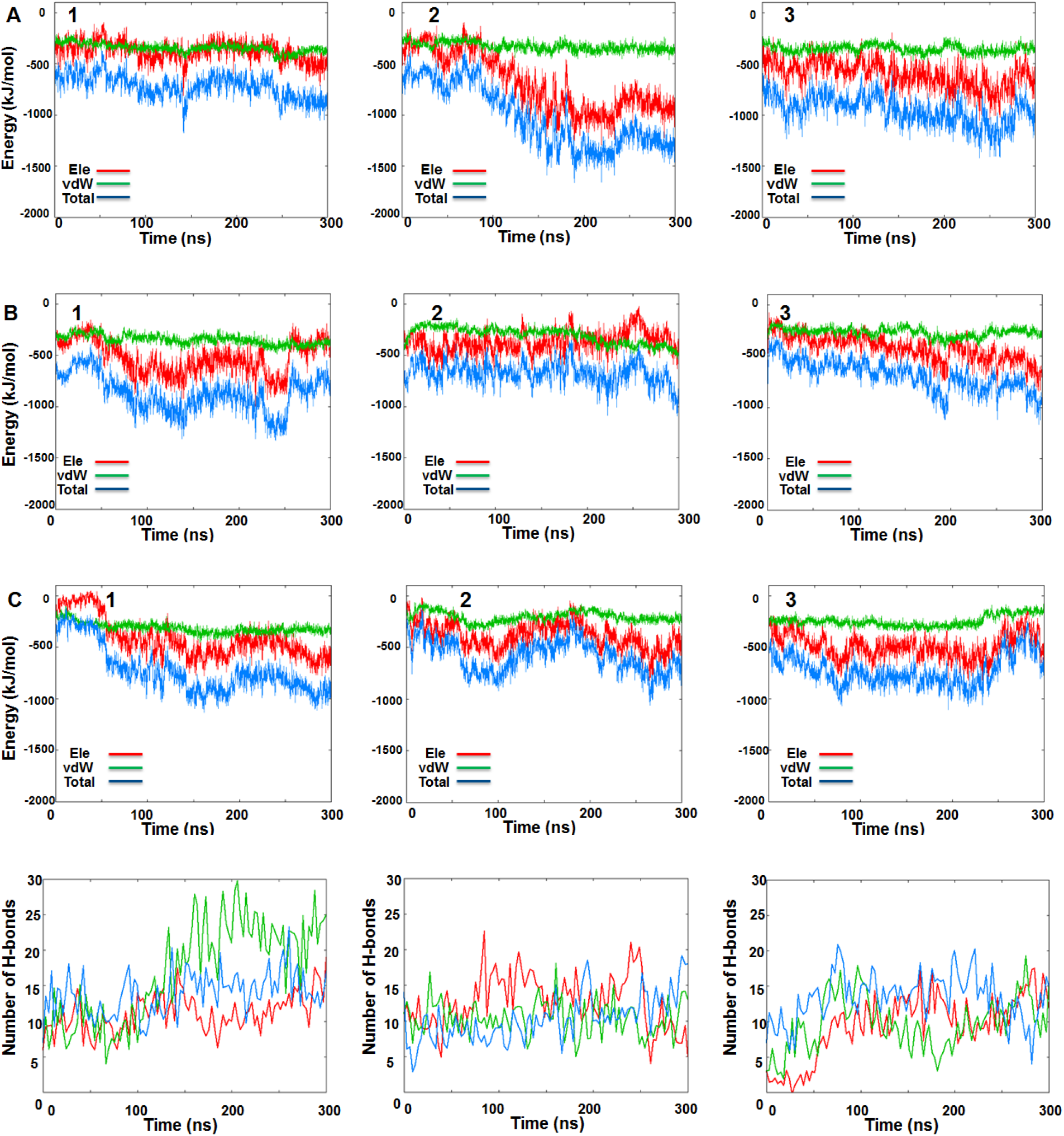
IRE1 face-to-face dimer MD simulations. Time-dependent interaction energy profiles for monomer A with monomer B during the three MD simulation replicas of (A) Native face-to-face crystal dimer (PDB code: 3P23), (B) KIRA-docked face-to-face dimer (PDB code: 3P23), (C) protein-protein docked pose of PDB 4U6R in face-to-face dimer form. Hydrogen bond analysis between monomers A and B during three MD replicas for (D) Native face-to-face crystal dimer (PDB code: 3P23), (E) KIRA-docked face-to-face dimer (PDB code: 3P23), (F) protein-protein docked pose of PDB 4U6R in face-to-face dimer form.

Moreover, energetic analysis of KIRA and ADP in each kinase active site of the face-to-face dimer revealed a lower energetic stabilization of ADP compared to KIRA, with each ligand being able to interact favorably with the IRE1 active site pocket for the entire simulation time during all three replicas (Figure S5).

### 3.4. MD simulations analysis: influence of KIRA on the back-to-back dimer

To investigate impact of KIRA on the IRE1 back-to-back dimer, three different systems were also considered here, namely the native back-to-back dimer crystal structure (PDB code: 4YZC), the native dimer structure (PDB code: 4YZC) with KIRA docked, and the protein-protein docked pose of PDB 4U6R in back-to-back dimer form, respectively. The stabilities of the three systems were studied during three MD replicas, each replica being 300 ns in length.

As reported in section 3.2 to assess the structural stability of the dimer structures, RMSD values of each dimer and the RMSD for the IRE1 dimer interface (Figure 7 and S6, respectively) were calculated. For each replica the structure present in the minimized, equilibrated system was used as the reference point and RMSD values were analyzed as functions of simulation time. The three replicas for the native back-to-back crystal dimer (PDB code: 4YZC) revealed that the IRE1 dimer is stable, as evidenced by low and relatively constant RMSD values of the three independent trajectories (Figure 7A and S6A). In contrast to the face-to-face dimer systems investigated, the KIRA-docked back-to-back dimer (PDB code: 4YZC) and the protein-protein docked pose of PDB 4U6R in the back-to-back dimer form show similar structural stabilities as the native IRE1 back-to-back dimer (Figures 7B, C and S6B and C).

**Figure 7.**
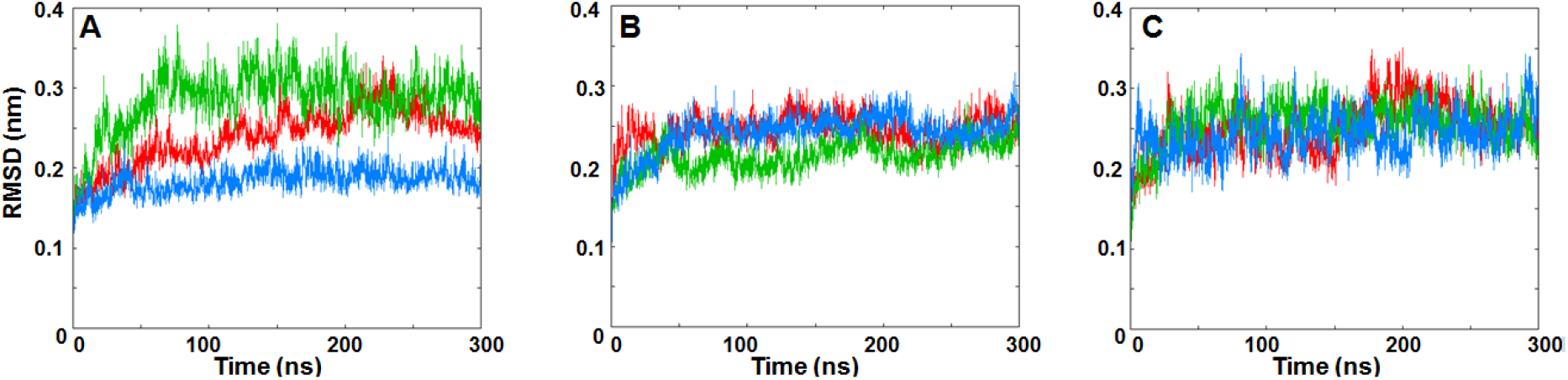
RMSDs of IRE1 back-to-back dimer Cα atoms during three MD replicas for (A) Native back-to-back dimer from crystal structure (PDB code: 4YZC), (B) KIRA docked in PDB 4YZC structure and (C) protein-protein docked pose of PDB 4U6R in back-to-back dimer form. Red for Replica 1, green for Replica 2 and blue for Replica 3, respectively.

Also for the three back-to-back systems, the interaction energy and the number of H-bonds between the monomers were analyzed (Figure 8). Energetic analysis of the occurring interchain interactions shows similar energetic stabilization of all three back-to-back dimers structures, and in all three replicas (Figure 8). The same trend was confirmed by the H-bonds analysis, with an equal number of H-bonds occurring between the native IRE1 back-to-back dimer compared to protein-protein docked pose and KIRA-docked back-to-back dimer (Figure 8). The overall analysis of these three IRE back-to-back dimers confirms our initial interpretation of the similar stabilities of the three dimer models. To further validate this hypothesis the last frame of each MD simulations were superposed with the native back-to-back dimer revealing an overall similar IRE1 back-to-back active conformation (Figure S7).

**Figure 8.**
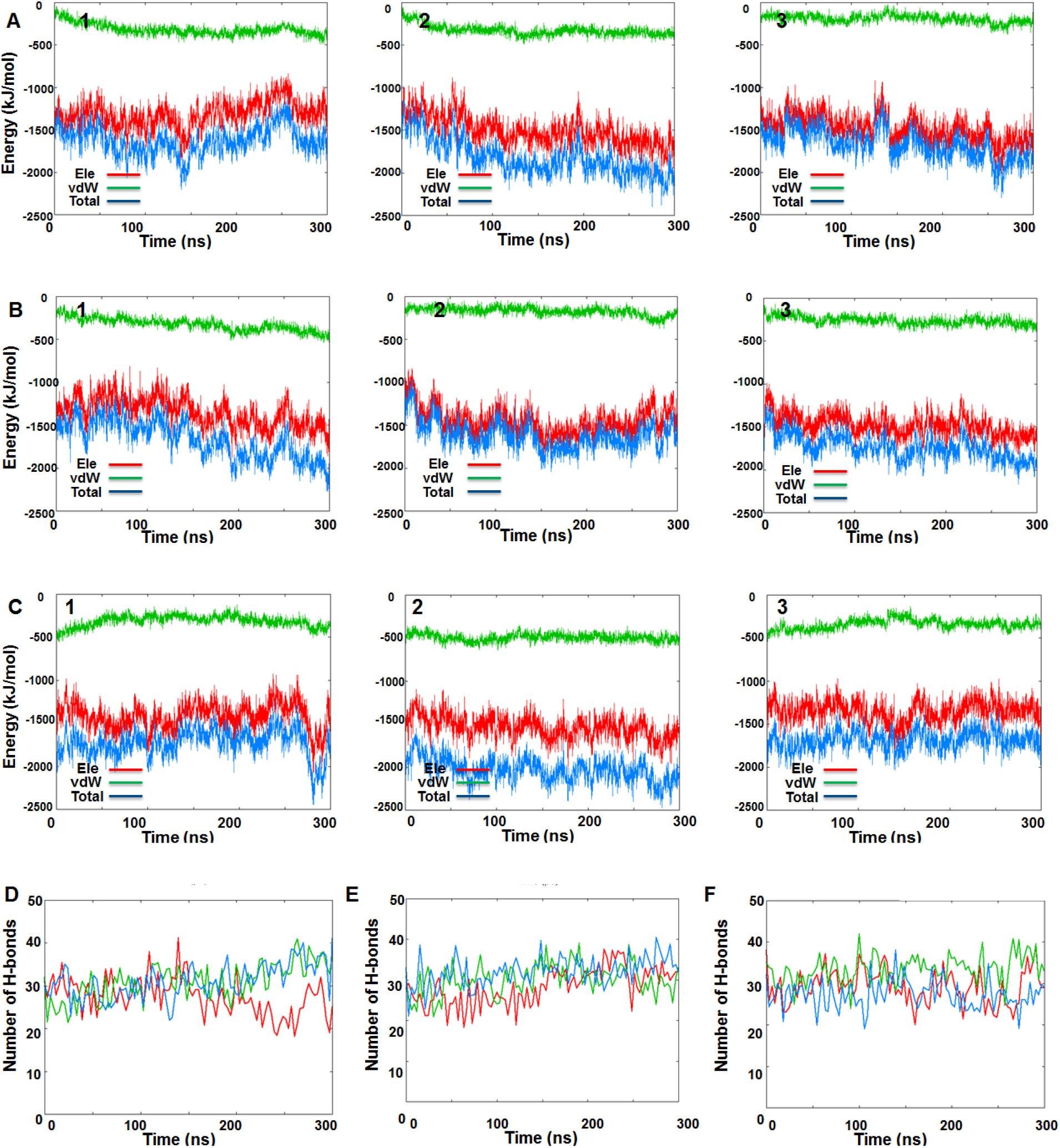
IRE1 back-to-back dimer MD simulation data for the three MD replicas. Time-dependent interaction energy profiles for monomer A with monomer B during the MD simulations of (A) native back-to-back dimer (PDB code: 4YZC), (B) KIRA docked in the 4YZC structure, (C) protein-protein docked pose of PDB 4U6R in back-to-back dimer. Hydrogen bond analysis between the monomers A and B during the three MD replicas for (D) native back-to-back crystal dimer (PDB code: 4YZC), (E) KIRA docked in PDB 4YZC structure, (F) protein-protein docked pose of PDB 4U6R in back-to-back dimer form.

Moreover, energetic analysis of KIRA and staurosporine in each kinase active site of the back-to-back dimer revealed similar energetic stabilization of staurosporine compared to KIRA, with each ligand being able to interact favorable with the IRE1 pocket active site during the entire simulation in the three replicas (Figure S8).

## 4. Conclusions and perspective

We have investigated the impact of KIRA binding on IRE1 dimer structures. Unexpectedly, the docking and MD simulations studies reveal that KIRA can bind to the kinase pocket of IRE1 in both the native face-to-face and back-to-back forms. A detailed analysis of the IRE1 monomer-monomer interactions process for the face-to-face dimer in presence of KIRA revealed energetic destabilization, suggesting that the binding of KIRA is affecting the system already at the stage of face-to-face dimer formation. Given that IRE1 activation appears to be dependent on the close communication between the kinase and RNase domains, the data leads us to believe that KIRA has a prominent role at the early stage of IRE1 activation, by destabilizing face-to-face dimer formation. This will impair the trans-autophosphorylation process, and thus preventing IRE1 from reaching the RNase active back-to-back structure.

The proposed mechanism of blocking the trans-autophosphorylation provides a molecular level validation of available experimental data where KIRA compounds inhibit IRE1 phosphorylation(8). This is further supported by the experimental observation that upon the inclusion of KIRAs, Western blotting reveals formation of IRE1 monomers only(8).

The data reported herein provide another small piece of information towards the understanding of IRE1 activity and the structural evidence of KIRA’s role in the IRE1 inhibition process, representing a stimulus to explore and better understand the IRE1 signaling.

## Acknowledgements

This research was funded by the EU’s Horizon 2020 research and innovation programme under the Marie Sklodowska-Curie grant 675448 (TRAINERS). The Faculty of Science at the University of Gothenburg and the Swedish Science Research Council (VR; grant number 2014-3914) are gratefully acknowledged for financial support (LAE) and the Swedish National Infrastructure for Computing for allocations of computing time at supercomputing centers C3SE and PDC.

## Supplementary material

RMSD data of top 5 docked poses (Table S1), Superposed structures displaying steric clashes (Figure S1), RMSDs of interface region in back-to-back dimers (Figure S2), and Structures at the three time points indicated in Figures 5D-F (Figure S3), are available as Supplementary material.

## Author contributions

All authors conceived the study. AC and CC performed the computations, analysed the data and wrote first draft. All authors revised the text.

## SUPPLEMENTARY INFORMATION

**Table S1.**
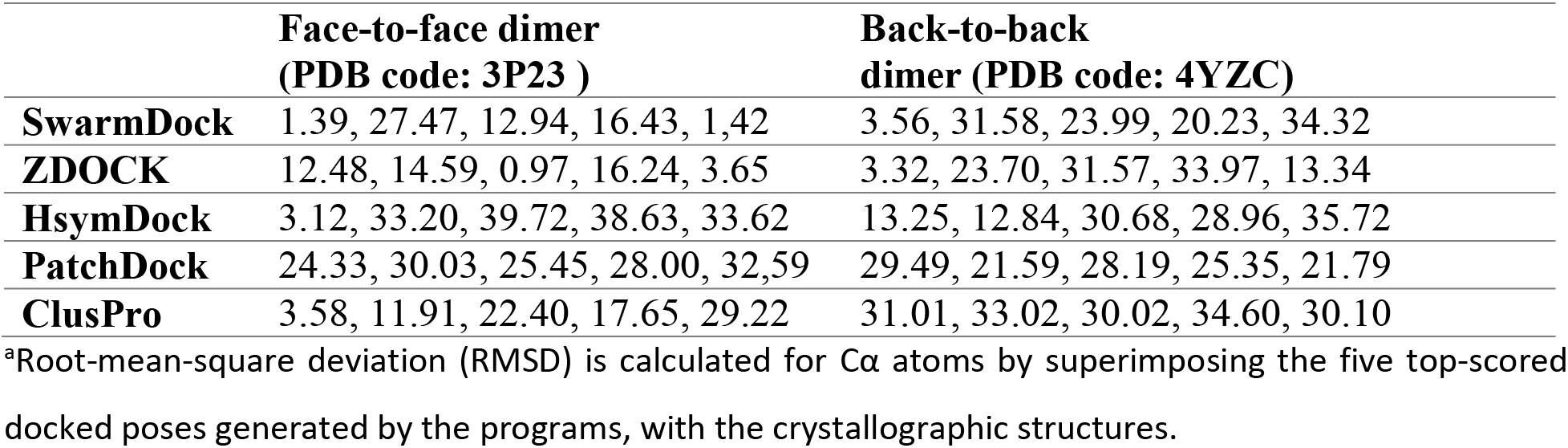
RMSD for the 5 top-scored docked poses generated using five different protein–protein docking approaches to reproduce the known IRE1 dimer complexes.

|  | **Face-to-face dimer**<br>**(PDB code: 3P23 )** | **Back-to-back**<br>**dimer (PDB code: 4YZC)** |
| --- | --- | --- |
| **SwarmDock** | 1.39, 27.47, 12.94, 16.43, 1.42 | 3.56, 31.58, 23.99, 20.23, 34.32 |
| **ZDOCK** | 12.48, 14.59, 0.97, 16.24, 3.65 | 3.32, 23.70, 31.57, 33.97, 13.34 |
| **HsymDock** | 3.12, 33.20, 39.72, 38.63, 33.62 | 13.25, 12.84, 30.68, 28.96, 35.72 |
| **PatchDock** | 24.33, 30.03, 25.45, 28.00, 32.59 | 29.49, 21.59, 28.19, 25.35, 21.79 |
| **ClusPro** | 3.58, 11.91, 22.40, 17.65, 29.22 | 31.01, 33.02, 30.02, 34.60, 30.10 |

<sup>a</sup>Root-mean-square deviation (RMSD) is calculated for Cα atoms by superimposing the five top-scored docked poses generated by the programs, with the crystallographic structures.

**Figure S1.**
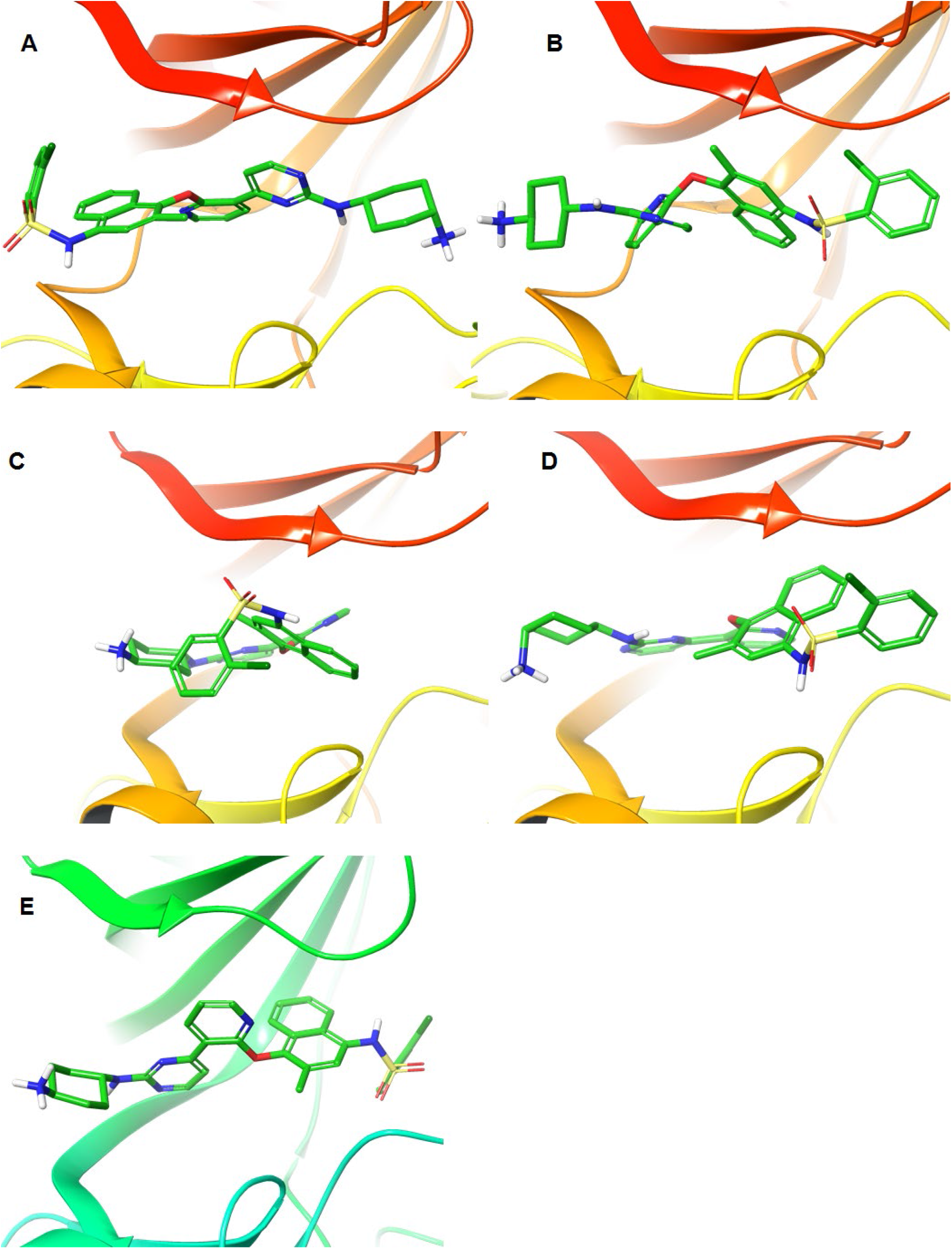
Representation of the KIRA docking pose in (A) Chain A of 3P23 PDB code, (B) Chain B of 3P23 PDB code, (C) Chain A of 4YZC PDB code, (D) Chain B of 4YZC PDB code. The crystallographic pose of KIRA in the 4U6R PDB structure is shown in panel E.

**Figure S2.**
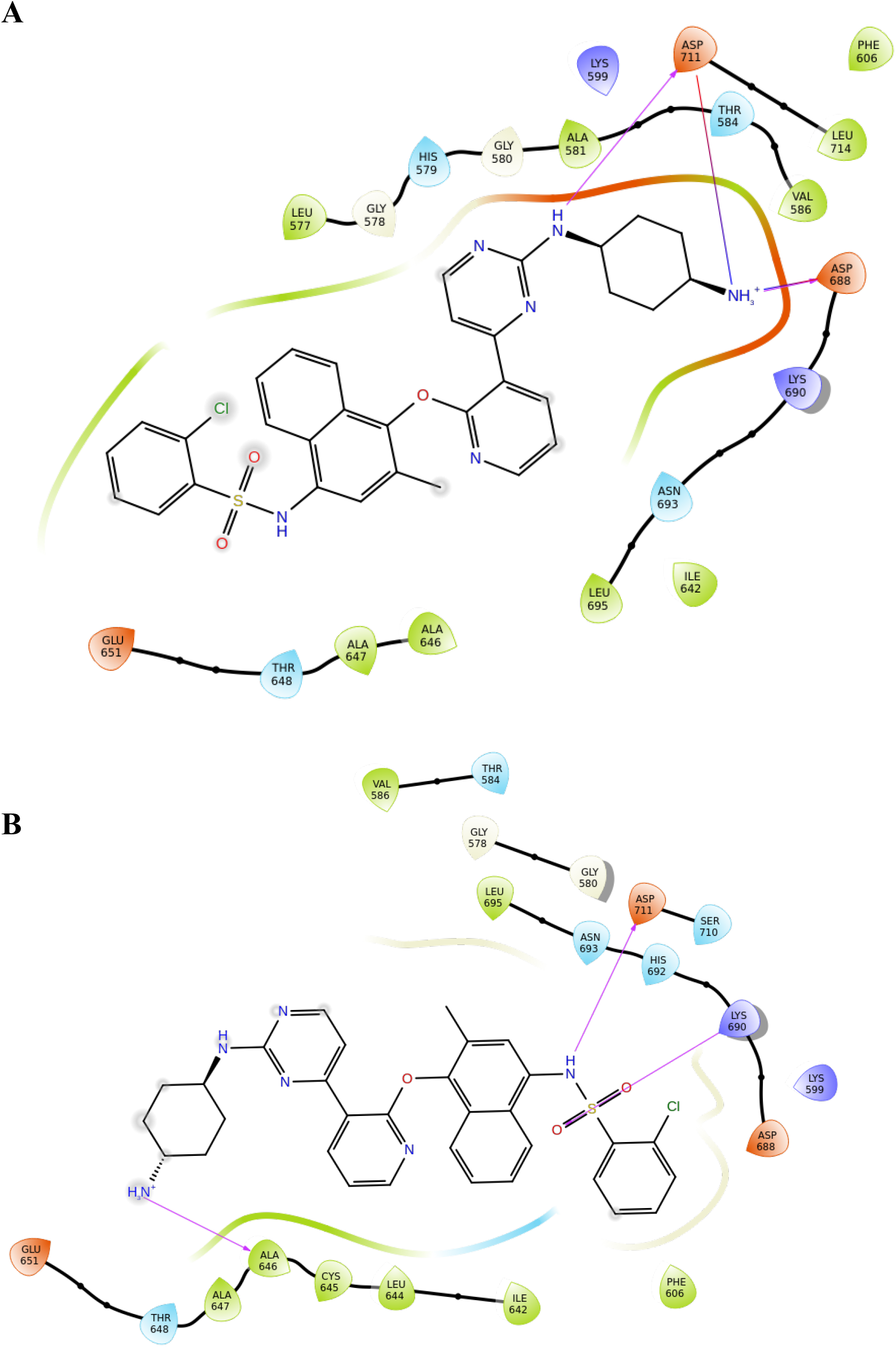
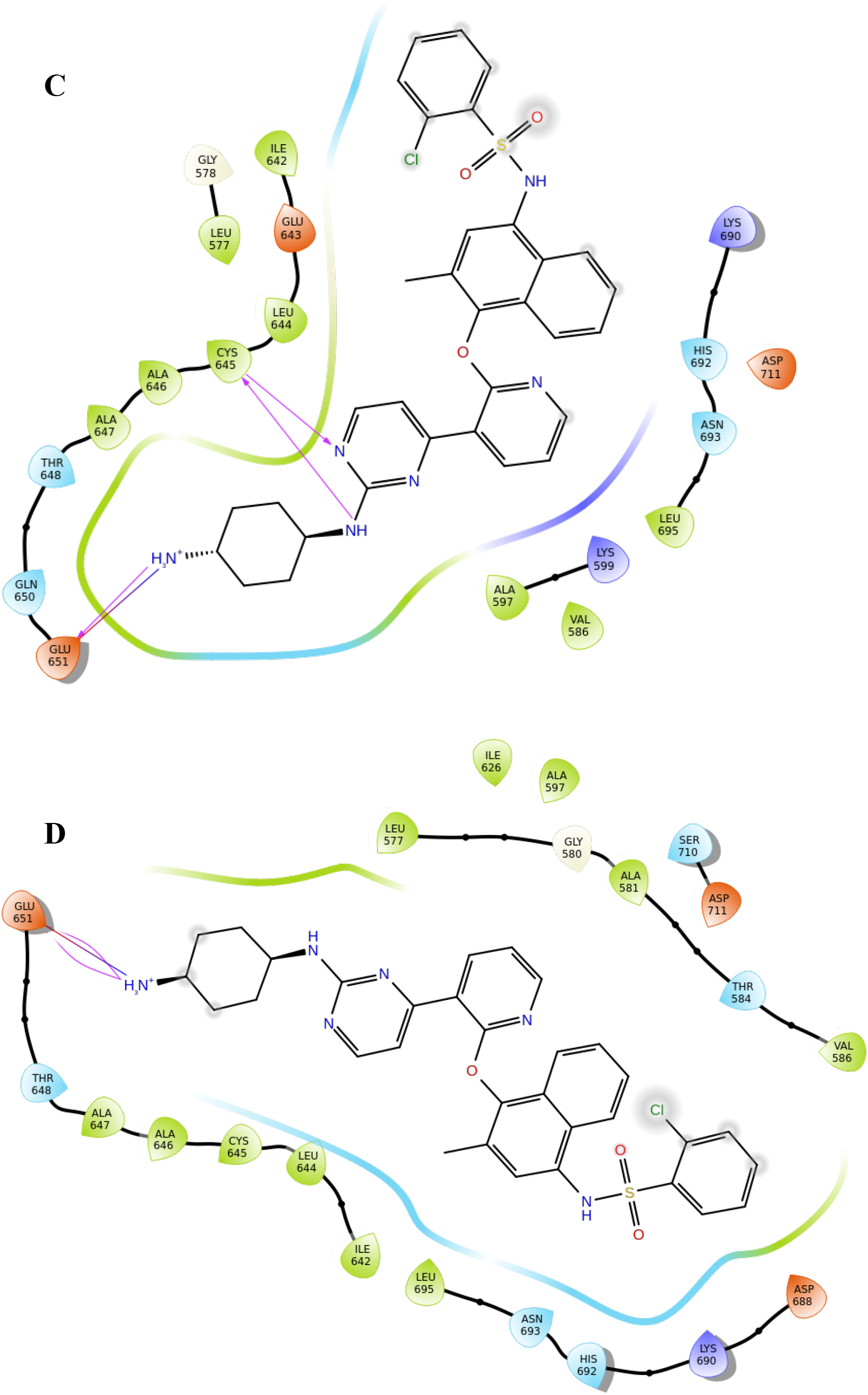
2D representation diagrams of the KIRA binding modes in (A) Chain A of 3P23 PDB code, (B) Chain B of 3P23 PDB code, (C) Chain A of 4YZC PDB code, and (D) Chain B of 4YZC PDB code.

**Figure S3.**
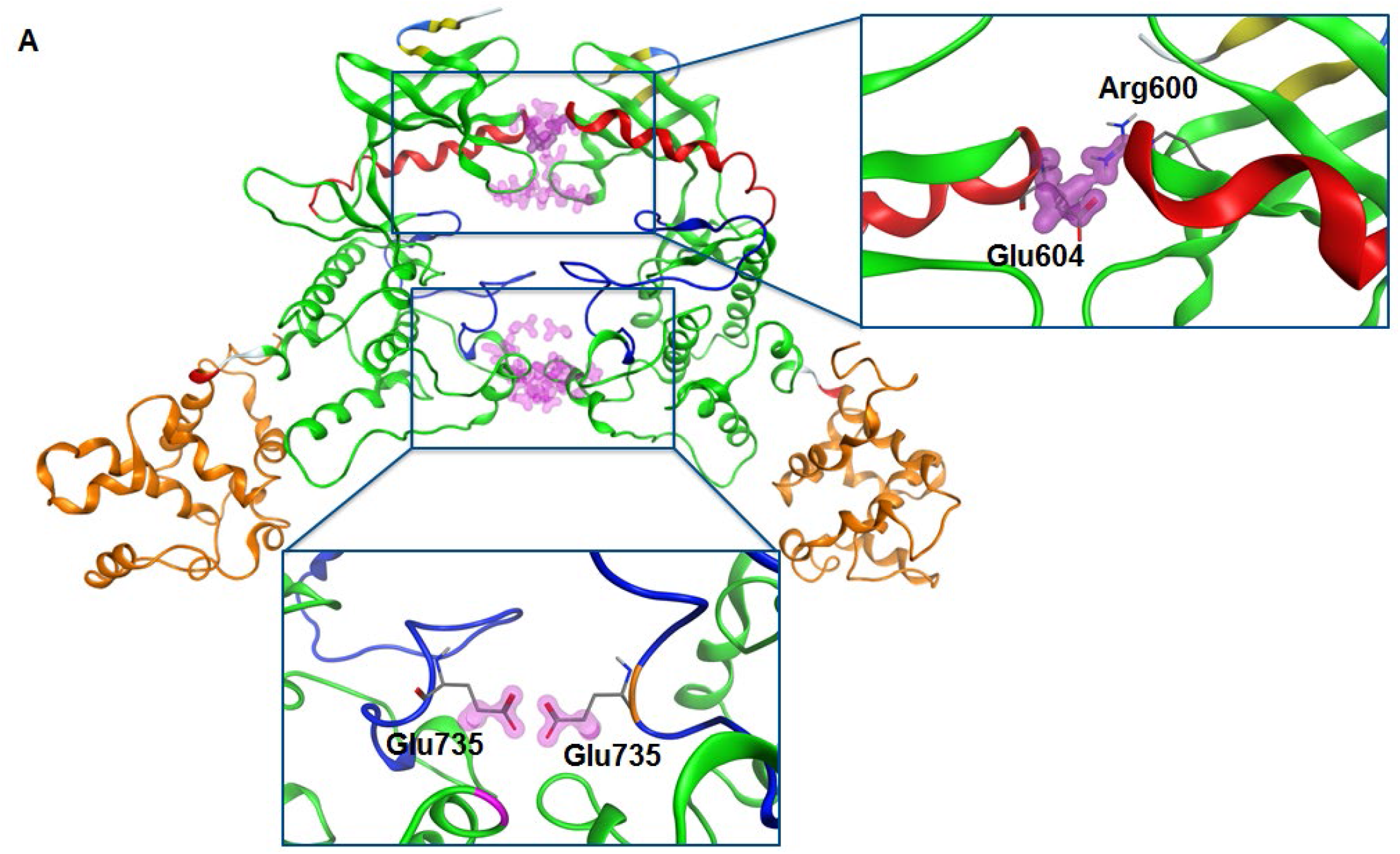
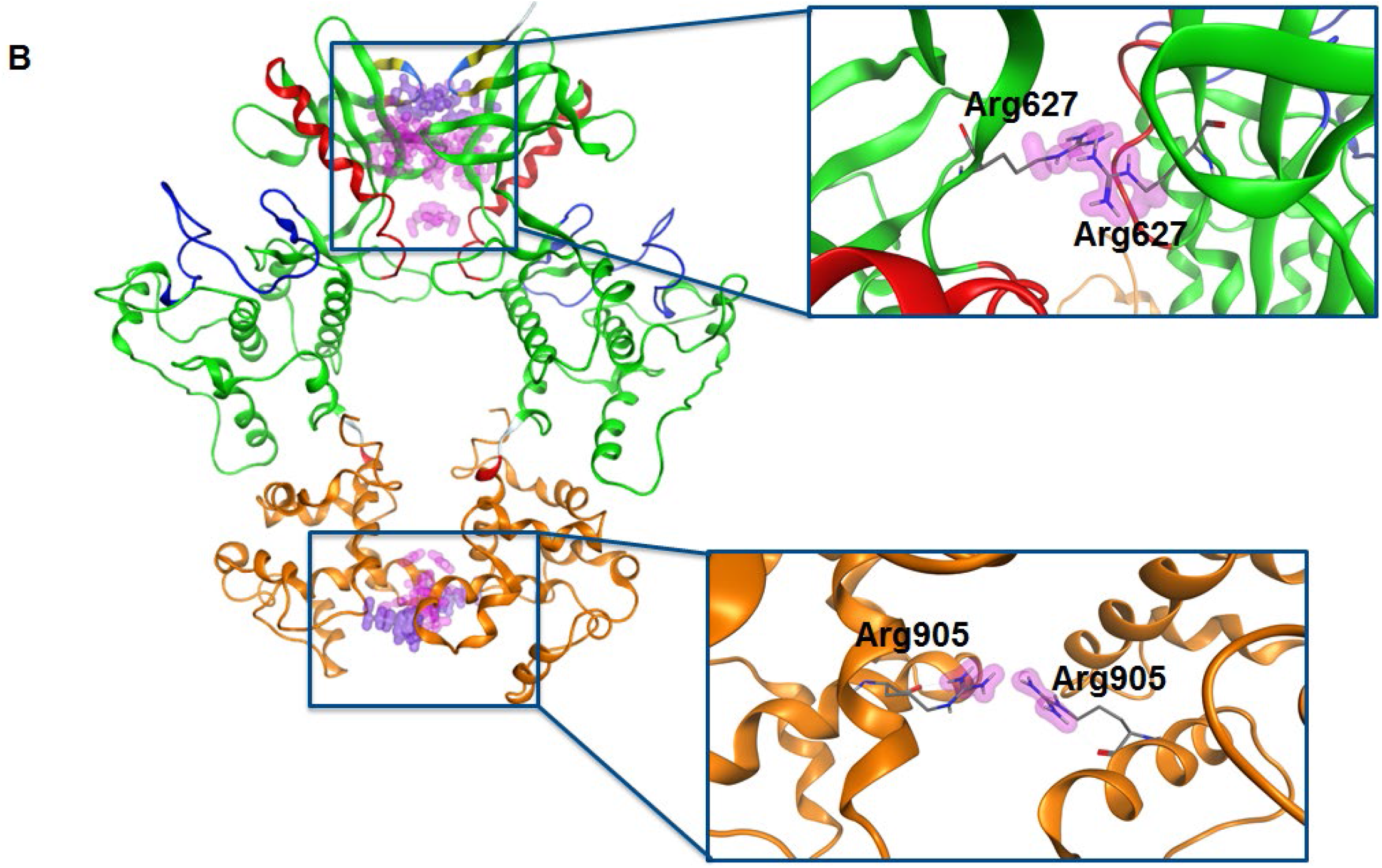
Ribbon diagram representing the structure of the KIRA-bound dimer forms obtained by superposition of the monomer of the 4U6R PDB structure on each monomer of the native crystallographic structures of the IRE1 in (A) face-to-face (PDB 3P23) and (B) back-to-back (PDB code: 4YZC) dimers. The kinase domain is shown in green (residues 571-832), the helix-αC in red (residues 603-623), the activation segment in blue (residues: 711-741) and the RNase domain in orange (residues 837-963). Violet spheres = steric clashes.

**Figure S4.**
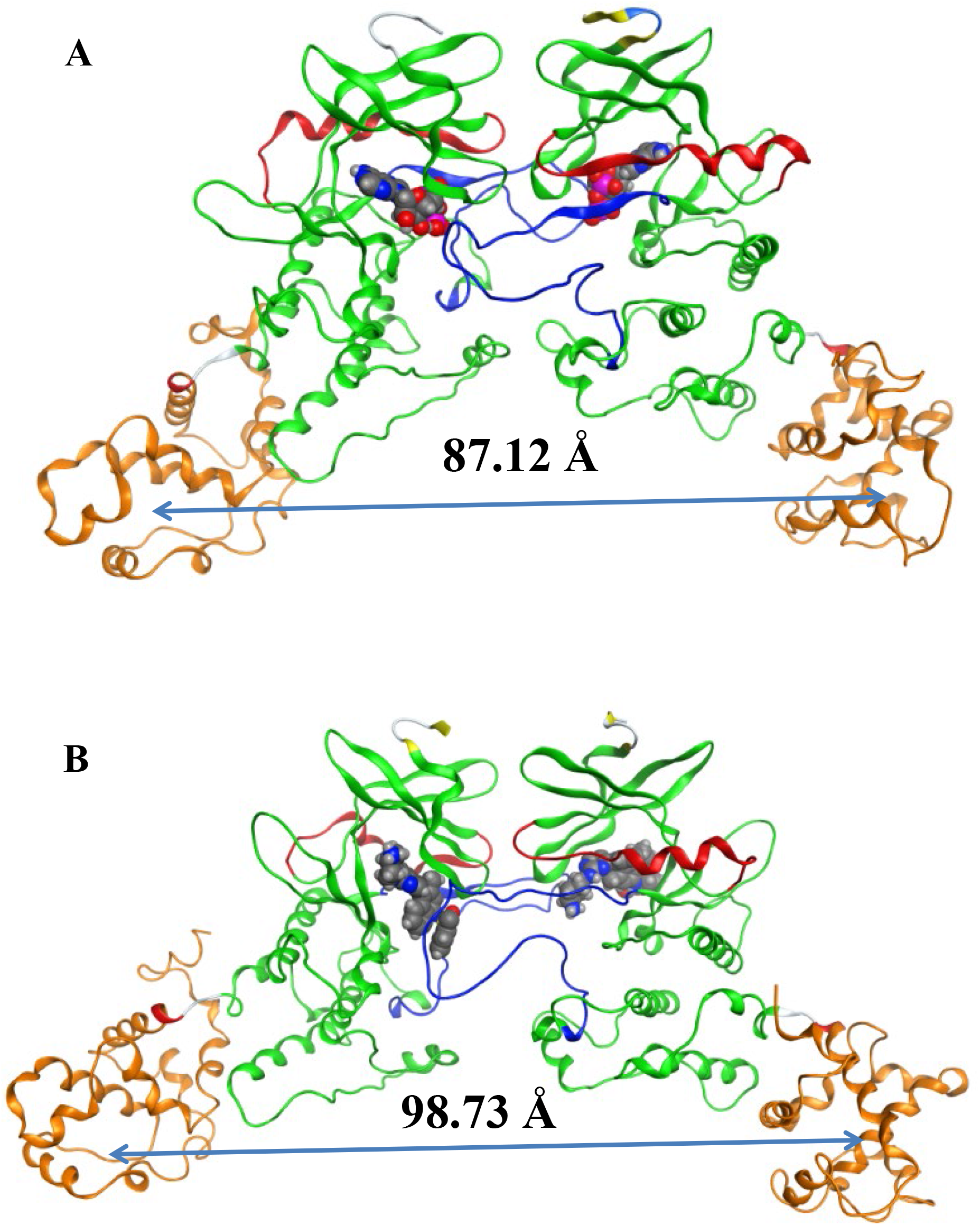
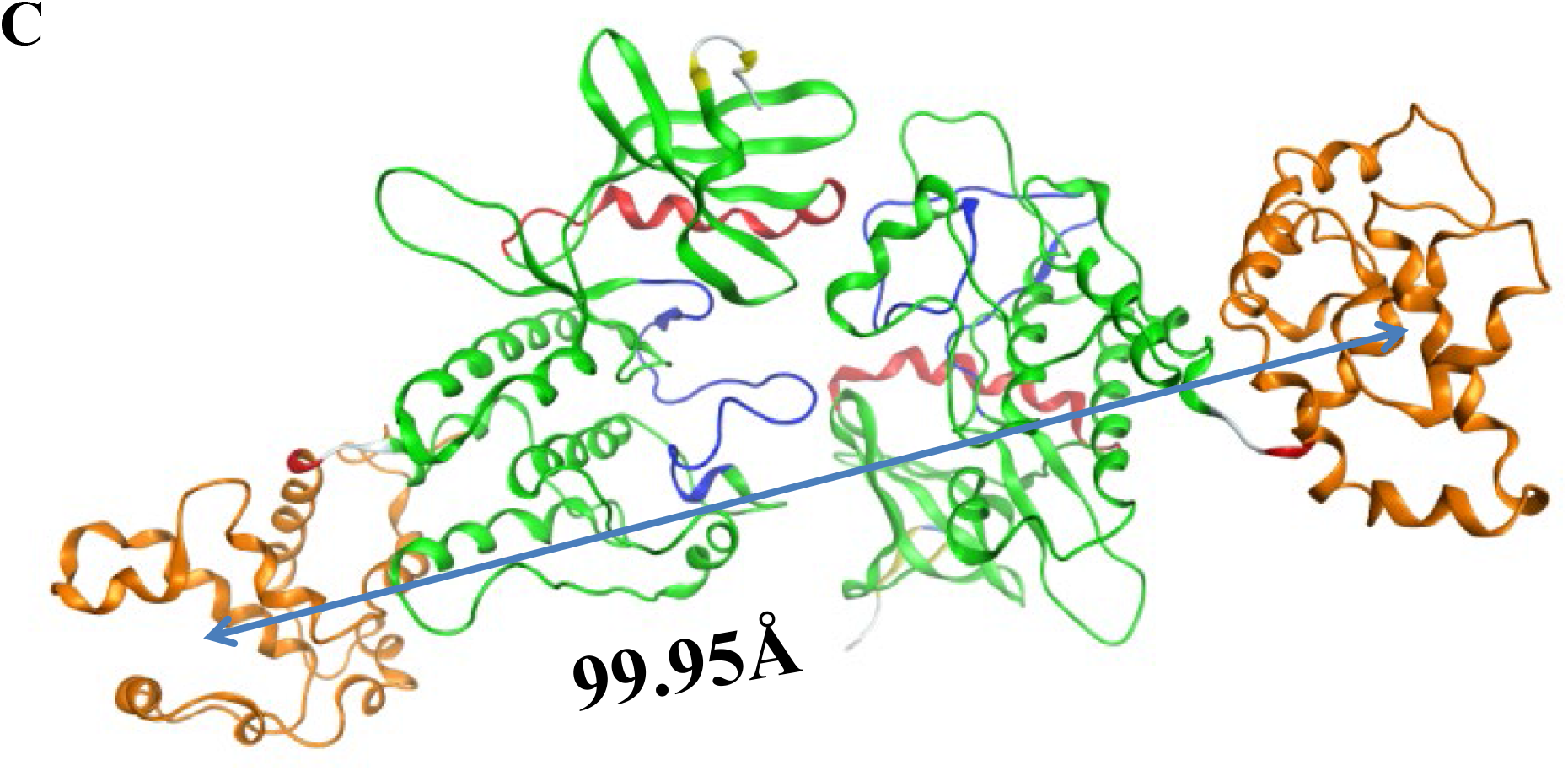
Comparison of the (A) native face-to-face crystal dimer structure (PDB code: 3P23) (point ‘a’ in Panel D of Figure 5), with the higher interface RMSD frame identified from three independent MD replicas for (B) KIRA docked in PDB 3P23 dimer (point ‘b’ in Panel E of Figure 5) and (C) protein-protein docked pose of PDB 4U6R in face-to-face dimer (point ‘c’ in Panel F of Figure 5). The distance between the RNase domain Center of Mass (COM) of dimer is shown. The kinase domain is shown in green (residues 571-832), the helix-αC in red (residues 603-623), the activation segment in blue (residues: 711-741) and the RNase domain in orange (residues 837-963). ADP (A) and KIRA (B) highlighted in space-filling model to indicate the kinase binding site.

**Figure S5.**
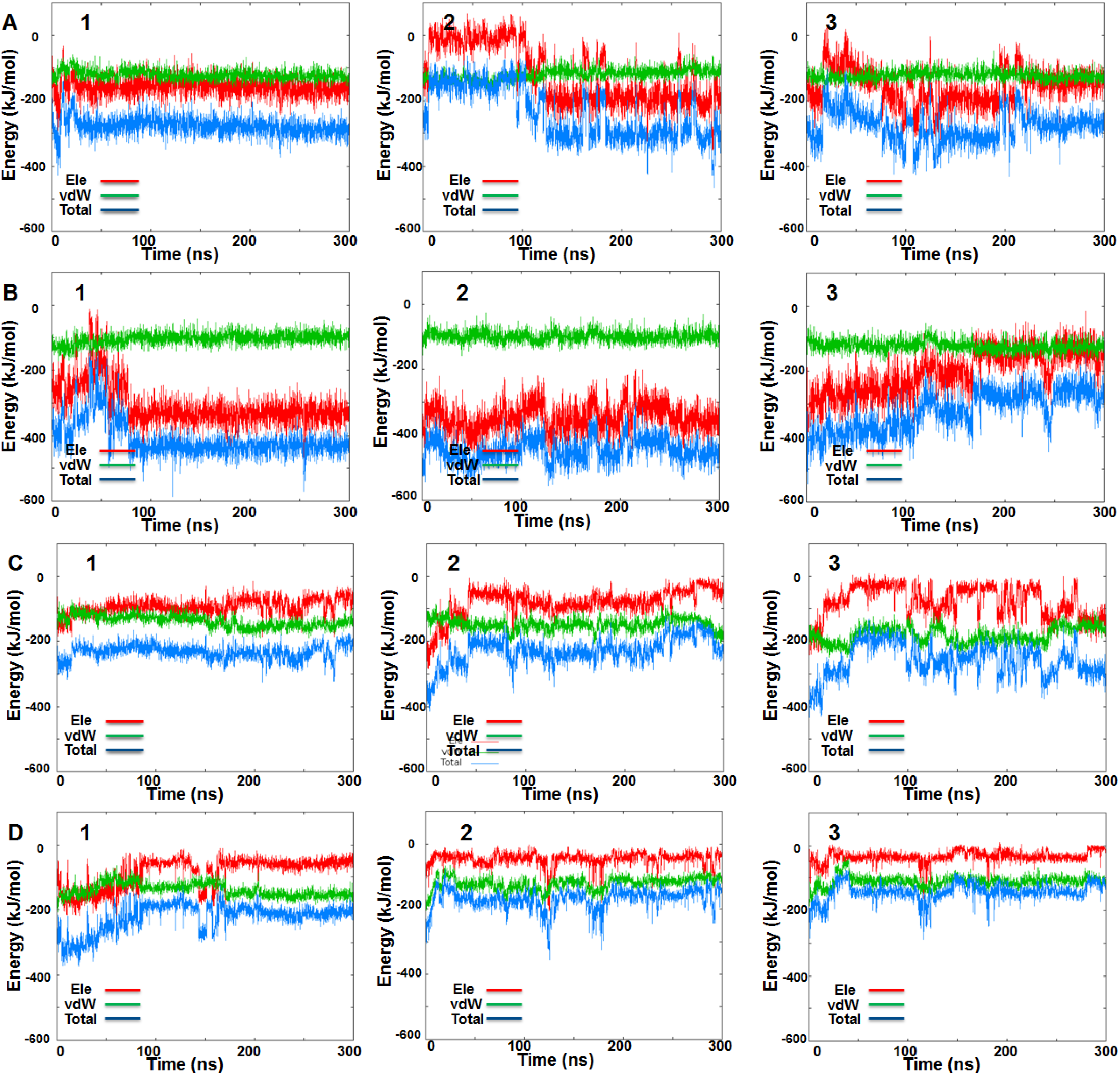
IRE1 face-to-face dimer MD simulations. Time-resolved interaction energy profiles for ADP during the three MD simulation replicas of the native face-to-face crystal dimer (PDB code: 3P23): (A) Chain A, (B) Chain B. Time-resolved interaction energy profiles during the three MD simulation replicas for KIRA docked in the native face-to-face crystal dimer (PDB code: 3P23): (C) Chain A and (D) Chain B.

**Figure S6.**
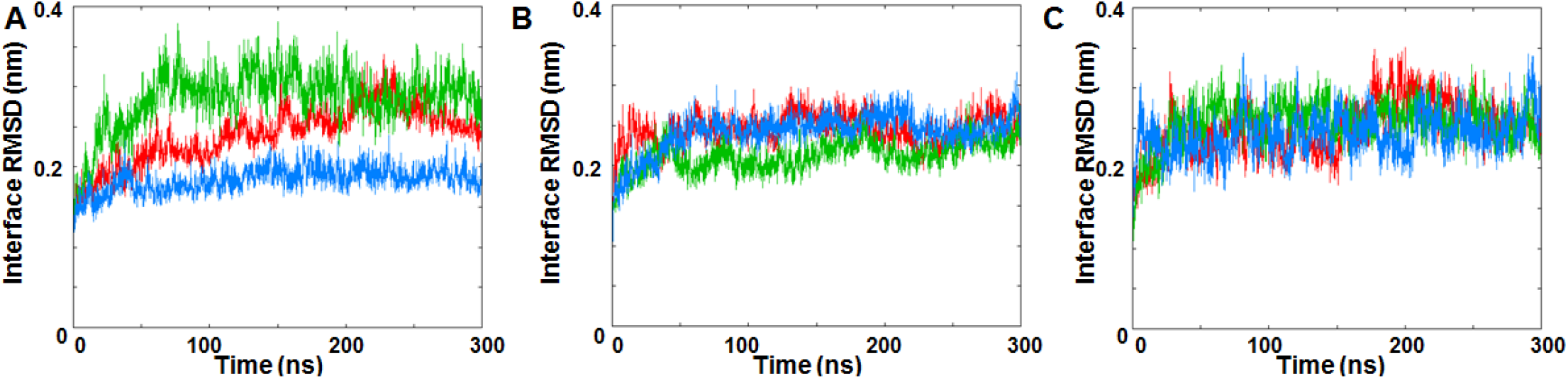
Interface RMSDs of IRE1 back-to-back dimer Cα atoms during the three MD simulation replicas of (A) native back-to-back crystal dimer structure (PDB code: 4YZC), (B) KIRA docked in PDB 4YZC dimer, and (C) protein-protein docked pose of PDB 4U6R in back-to-back dimer form. Replicates 1, 2, and 3 are represented in red, green and blue, respectively.

**Figure S7.**
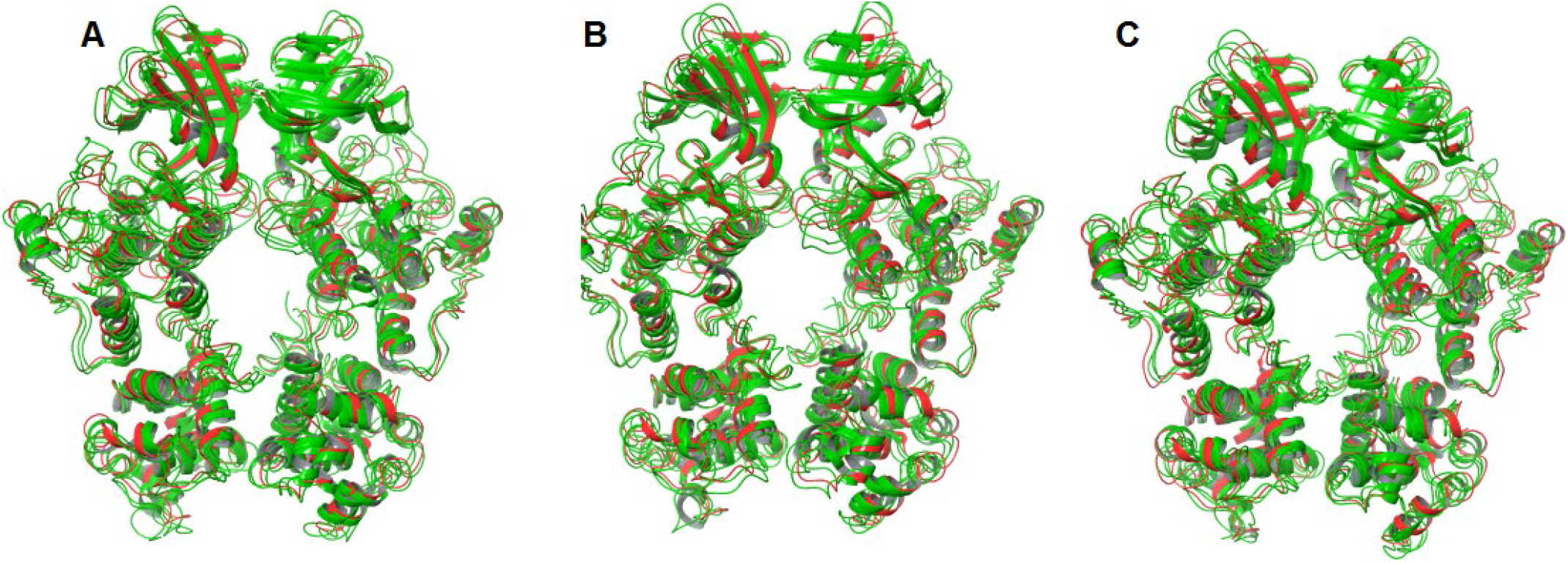
Superposition of the last frame of each individual MD simulation (green) of (A) native back-to-back dimer from crystal structure (PDB code: 4YZC), (B) KIRA docked in PDB 4YZC structure and (C) protein-protein docked pose of PDB 4U6R in back-to-back dimer, onto the native back-to-back crystallographic structure (PDB code: 4YZC) (red).

**Figure S8.**
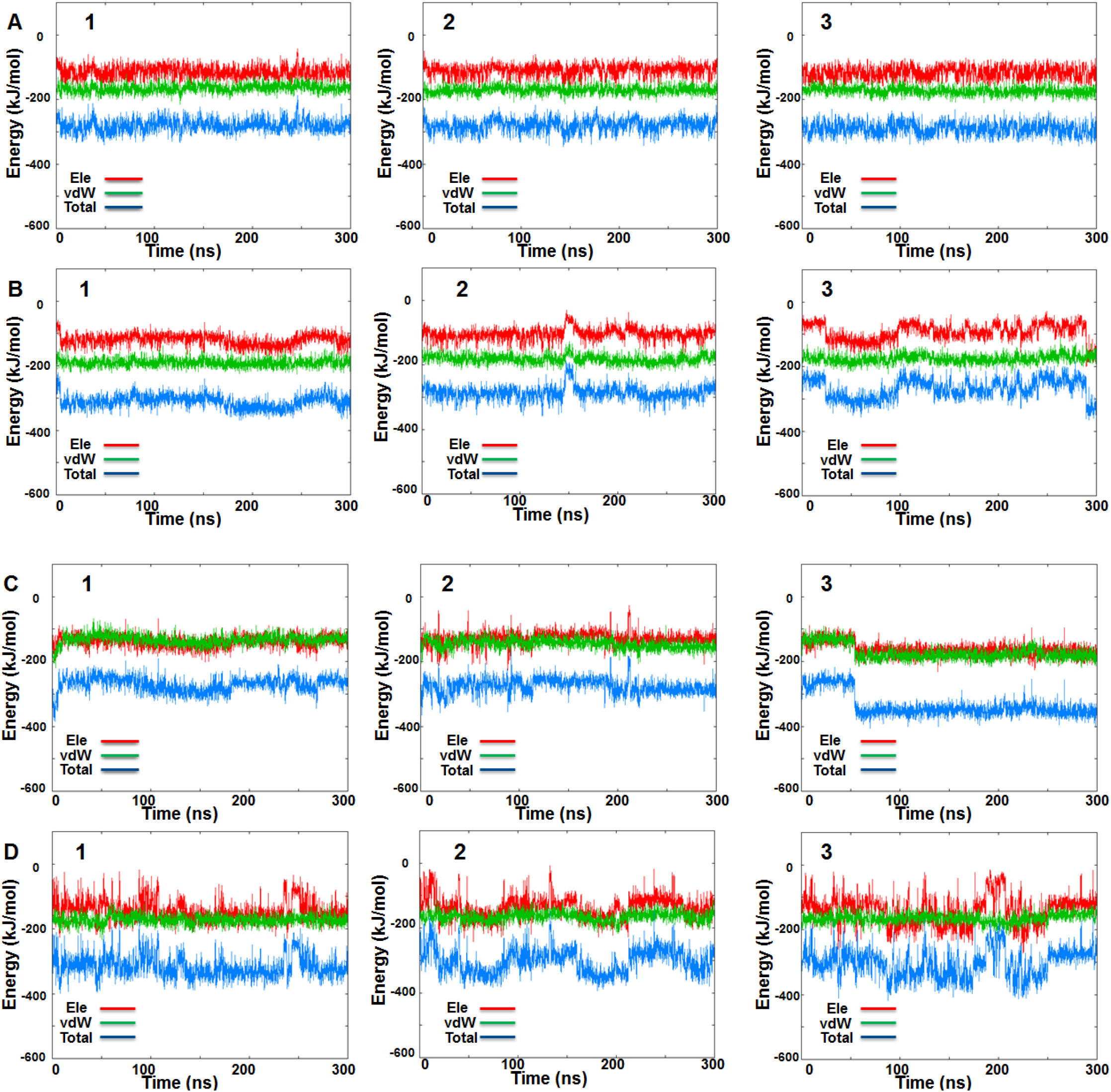
IRE1 back-to-back dimer MD simulations. Time-resolved interaction energy profiles for staurosporine during the three MD simulation replicas of the native back-to-back crystal dimer (PDB code: 4YZC): (A) Chain A, (B) Chain B. Time-resolved interaction energy profiles during the three MD simulation replicas for KIRA docked in the native back-to-back crystal dimer (PDB code: 4YZC): (C) Chain A and (D) Chain B.

